# Allelopathic effects of *Epichloë* fungal endophytes: experiment and meta-analysis

**DOI:** 10.1101/2021.11.10.468080

**Authors:** Heather A. Hager, Maija Gailis, Jonathan A. Newman

## Abstract

Host-specific *Epichloë spp*.endophytic fungal symbionts of pooid grasses that produce herbivore-deterring alkaloids and alter the grass host’s metabolite and protein profiles. Early observations suggested that *Epichloë* may have negative allelopathic effects on neighbouring plant species, particularly *Trifolium spp*. clovers, but subsequent allelopathy tests produced variable results. We examined two hypotheses: (1) *Epichloë* strains differ in allelopathic effect, and (2) *Epichloë* allelopathy negatively affects other plant species. We performed a greenhouse experiment using root exudates from Lolium perenne L. hosting different *E. festucae* var. *lolii* (Latch, M.J. Chr. & Samuels) C.W. Bacon & Schardl strains to compare their allelopathic effects on native legumes and forbs. We then used meta-analysis to examine the evidence to date for allelopathic effects of *Epichloë* endophytes. We found little difference in effect among *E. festucae* var. *lolii* strains and very little evidence for negative allelopathic effects of *Epichloë* in cool-season grasses across a range of methodologies, target plant species, and response measures. Negative allelopathic effects were detected only for root hair measures, which were from a single study. Positive effects on biomass were found for some experimental subgroups, including legumes. However, the majority of response variables showed no evidence for *Epichloë* allelopathy. Although there is currently little evidence for negative *Epichloë* allelopathic effects, our meta-analysis identified several research gaps. Experiments testing the functional belowground effects of *Epichloë* presence may help to determine its effects on non-host plant performance via plant-soil feedbacks.

## 1 Introduction

Plants release a variety of resource and non-nutritive compounds into the environment through root exudates, leachates, volatiles, and decomposing tissues. ^1–3^ Some of the non-nutritive compounds can have allelopathic effects, which may be positive or negative and are defined as stimulatory or inhibitory effects on the survival, growth, and functioning of other species. ^1,4,5^ Allelopathic plant-plant and plant-microbe interactions may be involved in structuring plant communities^6^ and facilitating the invasion of non-native plant species that release unfamiliar compounds in the invaded habitat.^6–8^ Many different types of allelopathic compounds have been isolated and identified in grasses, ^9,10^ although the specific modes of action in the affected organisms are often unknown. ^1,11,12, but^ ^see^ ^5^ In their recent systematic review of allelopathy research in grasslands, Silva *et al*.^13^ found that the overwhelming majority of studies reported negative allelopathic effects, indicating that allelopathy may be common in grassland ecosystems.

A number of cool-season pooid grasses have been introduced from their native ranges in southern Europe, northern Africa, and central Asia to new ranges in Oceania and the Americas for use in pasture, forage, and turf applications. Two such grasses, *Lolium perenne* (perennial ryegrass) and *Schedonorus arundinaceus* (tall fescue), have a long history of use in New Zealand and USA, respectively.^14–16^ Although these grasses are valued for maintaining productive and persistent pastures in their introduced ranges, livestock consuming them can experience a variety of health issues such as neurological and muscular disorders, decreased weight gain, and reproductive and thermoregulatory difficulties on some pastures but not on others. ^17–19^ These illnesses are related to the presence of species-specific symbiotic fungal endophytes of the genus *Epichloë* (formerly *Neotyphodium*) in the aboveground tissues of asymptomatic grasses: *E. festucae* var. *lolii* in *L. perenne* and *E. coenophiala* in *S. arundinaceus*. The fungi produce a number of herbivore-deterrent alkaloids, some of which are associated with livestock illnesses and others with herbivorous insect and nematode deterrence.^17,20–23^

Shortly after livestock illnesses were linked to the presence of *Epichloë* endophytes in the host grasses in USA, ^19^ researchers in New Zealand were noting reductions in legume abundance (typically *Trifolium spp*. clovers, often *T. repens*) in pastures where legumes were co-sown with endophyte-infected *L. perenne*. They suggested that legume decreases might be linked to alkaloids or other allelochemicals produced by the *Epichloë* symbiont or induced in the host grass. ^18,24–27^ Initial published laboratory assays of *Epichloë*-related allelopathy in *L. perenne* had mixed results, however, ranging from negative or no effects to positive allelopathic effects of the endophyte.^24,28–31^ Subsequent examinations of *Epichloë*-related allelopathy in *L. perenne* and other species also varied in their detection of allelopathic effects,^e.g., 32–34^ although some variation could be due to differences in study methodologies (e.g., target plant species, response variables measured). Some evidence suggests that *Epichloë* effects on target plants may be indirect, via soil microbial communities;^e.g., 35,36^ allelopathic compounds may alter microbe composition, and conversely, soil microbes may transform or moderate allelochemical effects. ^3,37^ Despite the diversity of results to date, *Epichloë*-related allelopathy continues to be suggested as an explanation for poorer plant performance in the presence of infected than uninfected host grasses and is used as an example of endophyte-driven allelopathy.^e.g., 38,39^

*Epichloë* species and strains differ in the main types of alkaloids they produce; ^40,41^ for example, the “common” *E. festucae* var. *lolii* strain, Lp19, produces ergovaline, peramine, and lolitrem B, ^42^ whereas that of *E. coenophiala* (unnamed) produces ergovaline, peramine, and lolines. ^43^ The discovery of ergovaline and lolitrem B as the main causes of livestock toxicity led to the search for *Epichloë* strains that produce only invertebrate-deterring alkaloids, and various *E. festucae* var. *lolii* and *E. coenophiala* strains are now available commercially in their host grass cultivars. ^44–46^. Endophyte strain affects multiple characteristics in the host grass, including alkaloid profiles, ^42,47^ volatile organic compound quantities, ^48^ root exudate chemical composition, ^49,50^ metabolite profiles, ^51,52^ and protein production.^52^ Thus, host grasses from the same genetic background but with different *Epichloë* strains may differ in their allelopathic effects, if such effects occur.

We performed an experiment and a meta-analysis to examine the evidence for allelopathic effects of *Epichloë* endophytes. Using a greenhouse experiment, we examined the allelopathic effect of four *Epichloë* festucae var. lolii strains in *L. perenne* cv. Alto on native North American grassland forbs and legumes and the introduced legume *Trifolium repens*. We hypothesized that endophyte presence in the host grass would result in negative allelopathic effects on the target plants and that endophyte strains would differ in their effects. We predicted lower biomass in target plants watered with root exudates from *Epichloë*-containing hosts than from endophyte-free hosts, with differences among *Epichloë* strains. We then used meta-analysis to synthesize the evidence to date for allelopathic effects on target plants due to *Epichloë spp*. in cool-season host grasses. We expected that endophyte presence in the host grasses would correspond with negative allelopathic effects on target plants. We also evaluated the effects of methodological differences among studies, specifically: host-endophyte pair, target plant functional group and species, method of target plant exposure to potential allelopathic substances, and substrate microbial status (sterile or inoculated). In all cases, we isolated the endophyte effect by using data on the allelopathic effect of grass hosts from the same genetic background (cultivar) with and without the endophyte.

## 2 Methods

### 2.1 Root exudate collection

Seeds of *Lolium perenne* L. cv. Alto that were endophyte-free or infected with one of four *E. festucae* var. *lolii* (Latch, M.J. Chr. & Samuels) C.W. Bacon & Schardl endophyte strains (AR1, AR37, NEA2, or Lp19) were provided by Agriseeds (Christchurch, New Zealand). Endophyte presence in seeds was confirmed using an immunoblot assay (Phytoscreen seed endophyte detection kit; Agrinostics, Watkinsville, Georgia, USA). Seeds were individually germinated and grown in rockwool medium (Grodan, Roermond, The Netherlands) for six weeks and then transferred to ceramic pots. Plants were then grown hydroponically for several weeks under greenhouse conditions of ∼40% relative humidity, 23 °C, and 16:8 h light:dark cycle. Pots were replenished daily with an all-purpose high-nitrate nutrient solution (20-8-20) and adjusted as necessary to maintain pH = 6. Once sufficient root volume was obtained, we began collecting root exudate solution. Plants that had been growing in nutrient solution for at least 24 h were transferred into new pots containing deionized water for 24 h and then returned to pots containing nutrient solution. The deionized water containing root exudates was collected and filtered using 25-*μ*m mesh to remove particulates. The procedure was repeated over a period of 6 wk to collect sufficient exudate solution for the experiment. Exudate solutions from plants containing the same endophyte strain were pooled and stored at 4 °C until they were used in the experiment (∼72 L).

### 2.2 Experimental setup

We used 11 native and one introduced grassland plant species as target plants: six perennial forbs from the family Asteraceae (*Achillea millefolium*, *Coreopsis lanceolata*, *Echinacea purpurea*, *Gaillardia aristata*, *Kuhnia eupatorioides* [=*Brickellia*], and *Ratibida pinnata*) and six perennial legumes from the family Fabaceae (*Baptisia alba*, *Dalea purpurea*, *Desmodium canadense*, *Glycyrrhiza lepidota*, *Lupinus perennis*, and *Trifolium repens*). Seeds were obtained from Wildflower Farm (Coldwater, Ontario, Canada), Prairie Moon Nursery (Winona, Minnesota, USA), or Home Hardware (Ontario, Canada; *Trifolium* only). Prior to planting, seeds of two legume species (*Lupinus* and *Baptisia*) were cold-stratified for 10 days according to the supplier’s planting instructions. No rhizobacteria were applied to legumes, except for *Trifolium*, which came pretreated.

Healthy seeds (not discolored or visibly damaged) of each target plant species were potted into seedling trays containing untreated Sunshine #4 potting mix (Canadian Sphagnum peat moss, coarse perlite, and dolomitic limestone; Sun Gro Horticulture, Agawam, Massachusetts, USA) and germinated in the greenhouse. Greenhouse conditions were 16:8 h light:dark (supplemented for ≥ 500 *μ*mol m^2^ s^*−*1^), 23 °C, and ∼10% relative humidity. Once the first leaf had emerged, 25 seedlings of each species were repotted individually into 1.3 L (10 × 10 × 13 cm) pots filled with the same potting soil, and 5 replicate seedlings of each species were randomly assigned to each exudate source (endophyte strain).

Five blocks were established on the greenhouse bench to account for a potential light gradient, and the five exudate sources and 12 target species were randomized within blocks, for a total of 300 pots. Each pot received reverse osmosis water in the first three days after repotting. From the fourth day onward, each pot received 50 mL of exudate solution every second day for the duration of the experiment (24 applications total = 1.2 L/pot over 7 wk). Reverse osmosis water was supplemented on non-exudate days if needed to maintain moist but not wet soil. Two days after the final exudate application, target plants were harvested, separated into shoot and root biomass, and dried to constant mass at 55 °C.

### 2.3 Statistical analysis

We used nested analysis of variance to examine the effects of exudate source endophyte strain, target plant functional group, target plant species nested within functional group, and their interactions on shoot, root, and total biomass and shoot:root ratio. Block was treated as a random effect. Significant fixed effects were examined using post-hoc Tukey tests. We used preplanned contrasts to compare target plant growth in exudate applications from endophyte-free vs. endophyte-present *Lolium perenne* for all species combined and within functional groups. The legume *Glycyrrhiza* was excluded from analyses involving root and total biomass because of missing root biomass data, and one extreme outlier for Baptisia-AR1 shoot:root ratio was removed. Shoot biomass and shoot:root ratio were log10-transformed, and root and total biomass were Box-Cox transformed to improve normality of residuals and homogeneity of variance. Analyses were performed in JMP 15 (SAS Institute, Cary, USA).

### 2.4 Meta-analysis

To identify previous studies of *Epichloë* allelopathic effects on neighbouring plant species, we searched the Web of Science citation index (Clarivate Analytics, Philadelphia, USA) on 20 July 2020 for all times and databases using the topic search: `endophyt* AND (allelo* OR feedback) AND (exudate OR conditioning OR conditioned OR extract OR leach*)`, resulting in 63 records. Examination of their titles and abstracts resulted in 18 potentially relevant articles for which we read the full text and checked the literature cited for additional studies. Studies were excluded for the following reasons: they compared only cultivars containing the endophyte and did not include an endophyte-free plant control; they compared endophyte-present and endophyte-free plants of different cultivars (i.e., confounded endophyte and cultivar effects); they did not contain the data required to calculate effect sizes (i.e., mean, sample size, and some measure of variance); they were review papers.

From each study, we extracted information about the host plant–endophyte pair and endophyte strain, the target plant species and functional group, target plant exposure method (soil conditioning, root exudates, aqueous tissue extracts, dried plant tissue), and the status of the soil in which target plants were grown (sterilized, live microbial inoculation, or untreated potting soil or filter paper). Because these study characteristics may affect the occurrence and strength of allelopathic effects, we extracted separate means, variance estimates, and sample sizes for each combination. Data were obtained from published tables and figures^53^ or directly from authors. We also included our data from the greenhouse experiment. The complete experimental and meta-analytical data sets are archived at Scholar’s Portal Dataverse. ^54^ Effect sizes were calculated as the standardized mean difference (Hedges’ g) using the `escalc()` function in the `metafor` package^55^ in R version 4.0.2 (R Core Team 2020) such that positive and negative effect sizes indicate positive and negative allelopathic effects of endophyte presence on target plant species, respectively.

We fitted random-effects meta-analytical models for individual response variables using the `rma()` function in `metafor`. We used both the standard method and the Hartung-Knapp adjustment to the standard error, and we report the widest 95% confidence intervals. ^56^ For the three response variables with the most studies (Table 1), we also fitted multilevel models using the `rma.mv()` function in metafor. Effect sizes were nested in study, and both effects were treated as random to account for nonindependence of effect sizes within studies (e.g., due to similar experimental conditions, use of a single control endophyte-free comparison for multiple effects, use of least significant difference when variance was not reported). We note, however, that sample sizes < 10–30 at the highest cluster level (i.e., study) can inflate type I error rates (i.e., increase false positives ^57–60^). We compared the fit of multilevel models with and without each within-study and between-study variance component using log-likelihood-ratio tests^60,61^ to determine whether they explained significant variance, and we used the `mlm.variance.distribution()` function in `dmetar` ^61^ to calculate the distribution of variance among the random effects. Finally, for responses with ≤ 4 effect sizes, we examined both composite (random-effects models) and individual effects (Table 1) due to known modelling issues with “small-k” (i.e., few effect sizes) meta-analysis. ^62^

**Table 1:** Meta-analytical models fitted to each response variable. Bold font indicates which model results are shown in the text. All model results are provided in the Supplemental Materials (Online Resource 1). ^*a*^M = multilevel random- and mixed-effects models, RE = random- and mixed-effects models using standard method, RA = random- and mixed-effects models with Hartung-Knapp adjustment to the confidence interval, SM = small-k analysis (comparing composite effect estimates with individual effects). An asterisk indicates that model results differed on whether the effect estimate’s 95% confidence intervals encompassed 0; otherwise, model results were in agreement. For nodulation score, the effect estimate was significant (positive) for the RE analysis, but the fail-safe number was 1.

| Response | Effect sizes | Studies | Models<sup>a</sup> |
| --- | --- | --- | --- |
| Shoot biomass | 133 | 12 | **M**, RE, RA |
| Root biomass | 121 | 9 | **M**, RE, RA |
| Total biomass | 120 | 8 | **M**, RE, RA |
| Germination or emergence (%) | 58 | 5 | RE, **RA** |
| Shoot length | 50 | 3 | RE, **RA** |
| Root length | 39 | 5 | RE, **RA** |
| Leaves/plant | 20 | 2 | RE, **RA** |
| Root hair density | 15 | 1 | RE, **RA** |
| Root hair length | 15 | 1 | RE, **RA** |
| Nodulation score | 9 | 2 | *RE, **RA** |
| Floral biomass | 7 | 2 | **RE**, RA |
| Total arbuscules + vesicles | 4 | 2 | * **RE**, **RA**, **SM** |
| Arbuscule numbers | 4 | 2 | **RE**, **RA**, **SM** |
| Vesicle numbers | 4 | 2 | * **RE**, **RA**, **SM** |

All meta-analytical models were fitted using inverse variance weighting and restricted maximum likelihood estimation. Homogeneity of effect sizes and the proportion of heterogeneity not attributable to sampling error were evaluated using Cochran’s Q and I^2^, respectively. Responses that showed significant heterogeneity were examined further using mixed-effects meta-regression models and each of four categorical predictors (moderators: host-endophyte combination, exposure method, target plant functional group, and soil status) to determine their moderating effect on the endophyte allelopathic effect. Moderators were fitted individually due to multicollinearity. ^61^ All model results are provided in Supplemental Materials (Online Resource 1) for comparison.

To evaluate the potential for research bias, we calculated fail-safe numbers (the number of nonsignificant effect sizes that would be required to make the overall effect nonsignificant) at alpha of 0.05 using the Rosenberg method (Rosenberg 2005) in `metafor` for effects for which the 95% confidence intervals did not encompass zero. We note that research bias tests may be unreliable (inflated type I error) due to data nonindependence, and such methods have not yet been extended robustly to multilevel models with moderators.^60,63,64^

## 3 Results

### 3.1 Greenhouse experiment

The exudate source endophyte strain differentially affected shoot biomass depending on the target plant species and functional group (significant exudate × species[functional group] and exudate × functional group effects; Table 2). Forb shoot biomass did not differ with exudate source. In contrast, legume shoot biomass was greater with NEA2 than endophyte-free exudate (and greater with endophyte-present than endophyte-free exudate; significant contrast, Table 2). This effect was driven by the legume *Glycyrrhiza*, which was the only species that differed significantly among exudate sources and was highest with NEA2 and AR1 and lowest with endophyte-free exudate (Figure 1).

**Table 2:**
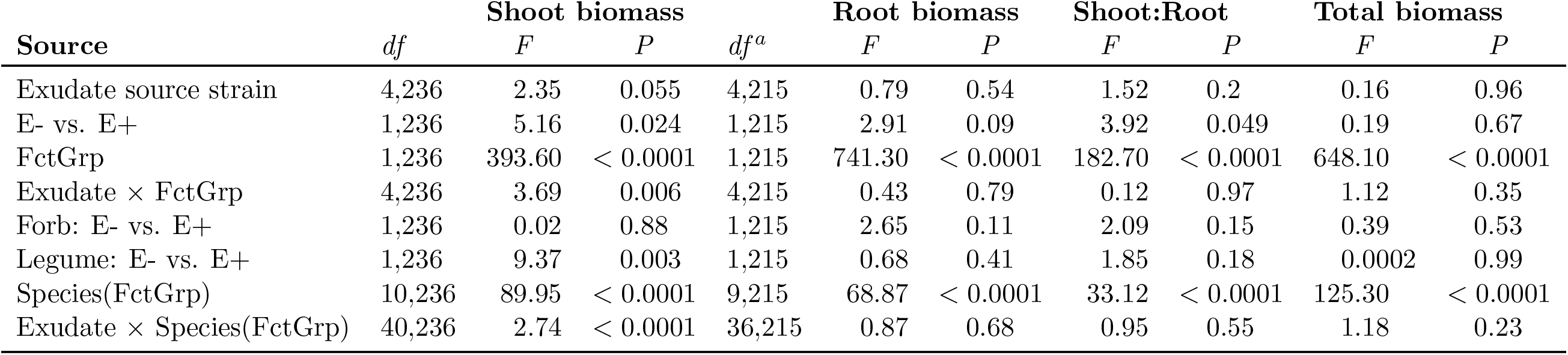
ANOVA results for effects of root exudates from Lolium perenne infected with different endophyte strains or no endophyte on growth responses of six native legume species (five legume species for measures involving root biomass) and six native forb species. Endophyte-free (E-) vs. endophyte-present (E+) contrasts are highlighted in grey. FctGrp = functional group (forb or legume). Highlighted rows are contrasts. Shoot biomass and shoot:root ratio were log10 transformed, and root and total biomass were Box-Cox transformed. ^*a*^Shoot:root *df* error = 214.

| Source | *df* | Shoot biomass |  |  | Root biomass |  | Shoot:Root |  | Total biomass |  |
| --- | --- | --- | --- | --- | --- | --- | --- | --- | --- | --- |
|  |  | *F* | *P* | *df*<sup>a</sup> | *F* | *P* | *F* | *P* | *F* | *P* |
| Exudate source strain | 4,236 | 2.35 | 0.055 | 4,215 | 0.79 | 0.54 | 1.52 | 0.2 | 0.16 | 0.96 |
| E- vs. E+ | 1,236 | 5.16 | 0.024 | 1,215 | 2.91 | 0.09 | 3.92 | 0.049 | 0.19 | 0.67 |
| FctGrp | 1,236 | 393.60 | < 0.0001 | 1,215 | 741.30 | < 0.0001 | 182.70 | < 0.0001 | 648.10 | < 0.0001 |
| Exudate × FctGrp | 4,236 | 3.69 | 0.006 | 4,215 | 0.43 | 0.79 | 0.12 | 0.97 | 1.12 | 0.35 |
| Forb: E- vs. E+ | 1,236 | 0.02 | 0.88 | 1,215 | 2.65 | 0.11 | 2.09 | 0.15 | 0.39 | 0.53 |
| Legume: E- vs. E+ | 1,236 | 9.37 | 0.003 | 1,215 | 0.68 | 0.41 | 1.85 | 0.18 | 0.0002 | 0.99 |
| Species(FctGrp) | 10,236 | 89.95 | < 0.0001 | 9,215 | 68.87 | < 0.0001 | 33.12 | < 0.0001 | 125.30 | < 0.0001 |
| Exudate × Species(FctGrp) | 40,236 | 2.74 | < 0.0001 | 36,215 | 0.87 | 0.68 | 0.95 | 0.55 | 1.18 | 0.23 |

**Figure 1:**
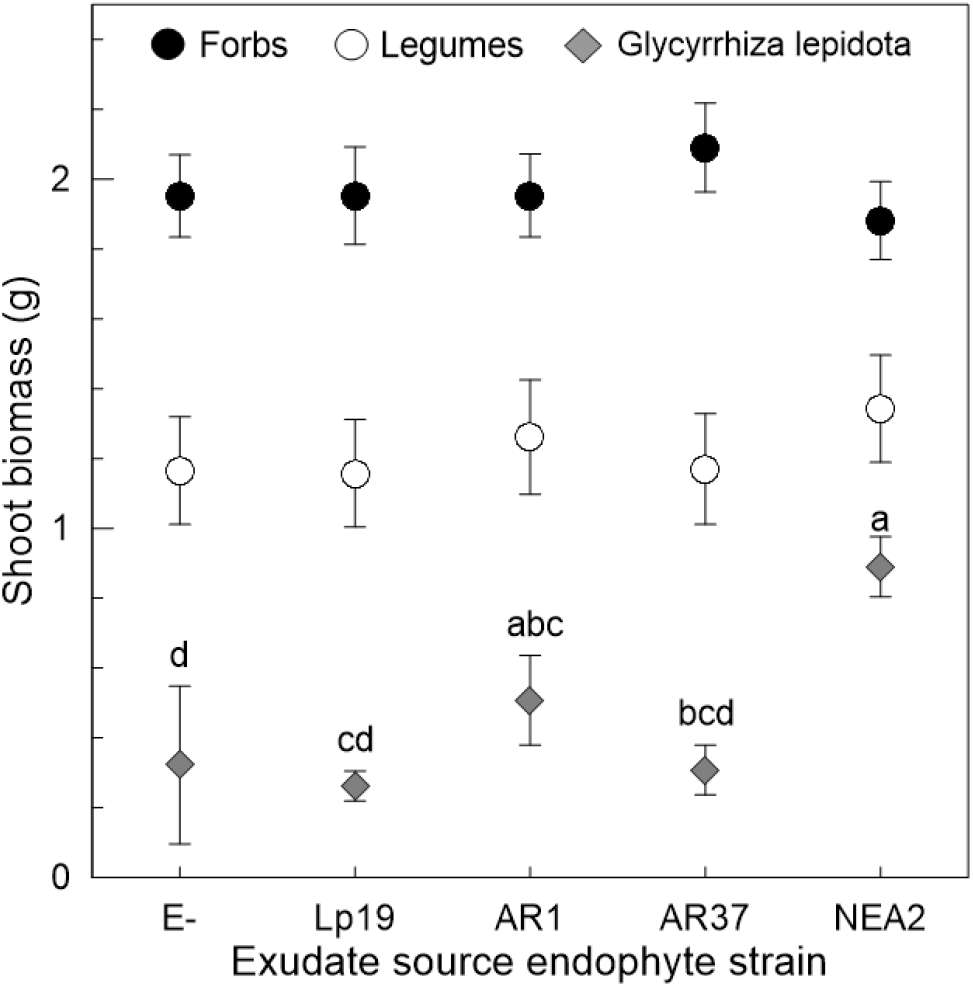
Mean ± standard deviation of untransformed shoot biomass of forbs, legumes, and the legume *Glycyrrhiza lepidota* watered with root exudates from *Lolium perenne* endophyte-free (E-) or infected with one of four *Epichloë festucae* var. *lolii* strains.

Root biomass, shoot:root ratio, and total biomass were mostly unaffected by the exudate source. Only shoot:root ratio was greater with endophyte-present than endophyte-free exudate sources (significant contrast), but the effect was very small. Unsurprisingly, all growth measures differed by target plant species and functional group (significant species[functional group] and functional group effects; Table 2). Forbs generally produced more shoot, root, and total biomass and had lower shoot:root ratios than legumes (not shown).

### 3.2 Meta-analysis

Twelve articles were both relevant and had data that could be extracted from figures or tables or obtained from the authors. ^24,30,32–36,65–69^ Four additional papers^28,29,31,70^ and one thesis^71^ identified from the references and our knowledge of the literature also had relevant extractable or obtainable data. Data from these 17 sources and our greenhouse experiment provided 603 effect sizes.

Data were obtained for 17 response variables. Two responses with one effect size each (stolons per plant and plant vigour score), and one response with two effect sizes (arbuscle:vesicle ratio), were excluded from further analysis. The remaining responses each had ≥ 4 effect sizes and were examined meta-analytically (599 effect sizes analysed). There were 41 target plant species: 12 legumes, including six *Trifolium* species; 11 forbs; nine trees; and nine grasses, including four of the host grasses (considered as selfE+ and selfE− functional groups when source and target plant species were the same, i.e., autoallelopathy, and the target plant did or did not host the endophyte, respectively). Four exposure methods were used: “aqueous extracts” obtained by soaking dried host-grass tissue in water; “dried host-plant tissue” applied directly on or in the soil; “root exudates” collected from the host grass; and “soil conditioning” by growing and removing the host-grass from soil that was then planted with the target plant. Finally, five host–endophyte pairs (all having strictly vertical endophyte transmission) were examined: *Festuca rubra*–*E. festucae* (48 effect sizes, one study), *Lolium multiflorum*–*E. occultans* (46 effect sizes, four studies), *Lolium perenne*–*E. festucae* var. *lolii* (287 effect sizes, eight studies), *Schedonorus arundinaceus* –*E. coenophiala* (205 effect sizes, five studies), and *Schedonorus pratensis*–*E. uncinata* (13 effect sizes, two studies).

Meta-analysis showed negative effects of *Epichloë* presence-related allelopathy on target plant root hair density and root hair length (Figure 2A). Both effects were large (> 0.8; Cohen 1988), with large fail-safe numbers, and the low I^2^ values (< 40%) suggest that effect size heterogeneity has minimal importance. ^72^ However, all effect sizes were from a single study. In contrast, there was a small (0.2) positive allelopathic effect on target plant shoot biomass (Figure 2A), with a large fail-safe number. Although the heterogeneity test was statistically significant (Q132 = 181.78, *P* = 0.003), likely due to the large number of effect sizes, the low I^2^ suggests that its importance is minimal.

**Figure 2:**
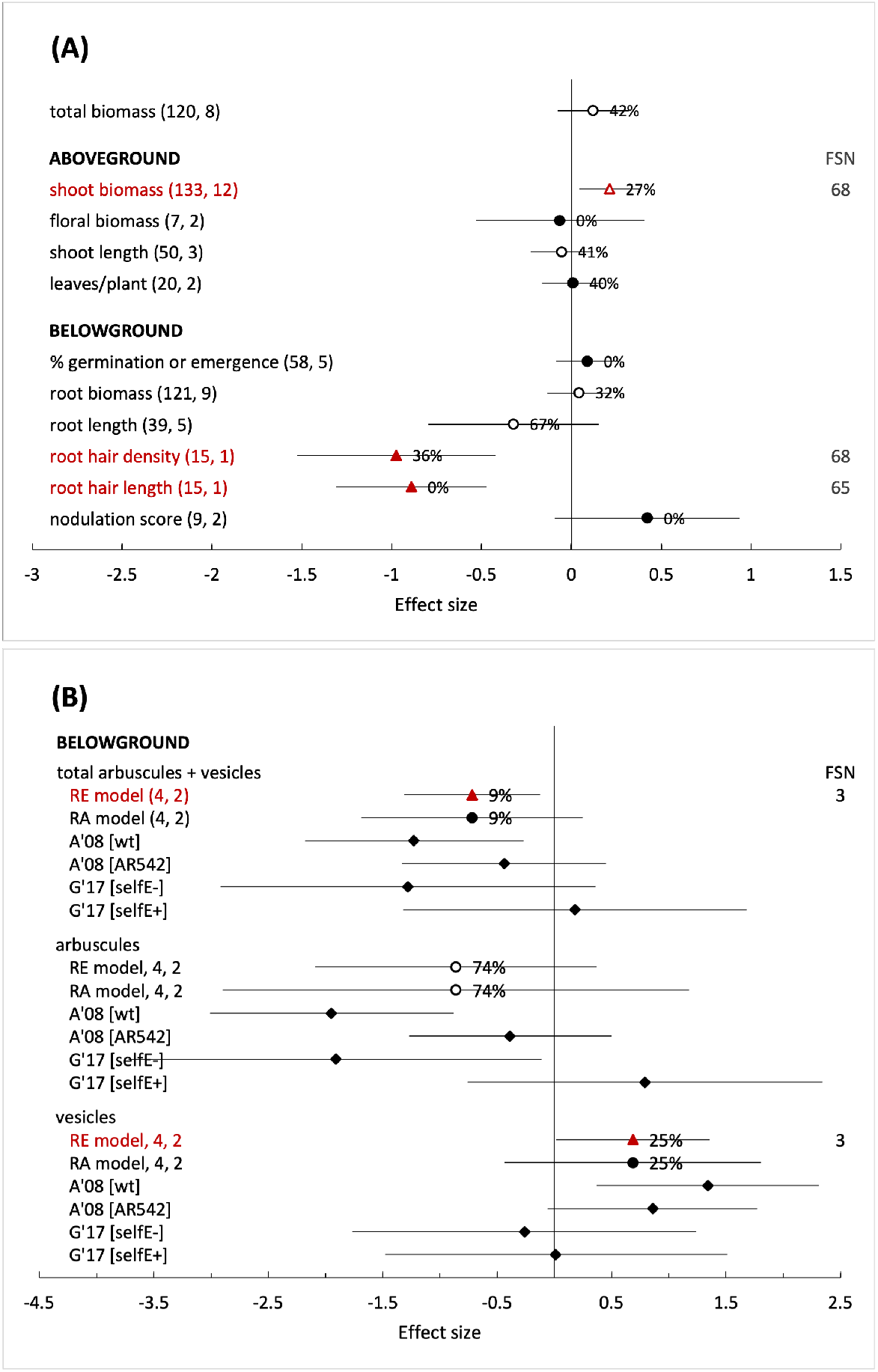
Overall target-plant responses to allelopathy due to the presence of *Epichloë* endophyte in its host grass. (A) Responses with 7–120 effect sizes. Points and whiskers are effect sizes and 95% confidence intervals, respectively. Numbers of effect sizes and studies are in parentheses. Point labels are I^2^ values (proportion of heterogeneity not attributable to sampling error). Dark red triangles = significant composite effects, circles = nonsignificant composite effects, hollow symbols = significant heterogeneity of effect sizes (Cochran’s Q). FSN = fail-safe numbers (number of nonsignificant effect sizes required to make the overall effect nonsignificant). (B) Responses with four effect sizes. Symbols as in (A) and diamonds = single effect sizes. A’08 = Antunes *et al*. 2008, ^35^ G’17 = Garcia-Parisi and Omacini 2017, ^65^ and brackets indicate treatment differences: wt and AR542 are different endophyte strains; selfE− and selfE+ indicate that target plants were the same species as the host grass and did not or did host the endophyte, respectively. See Table 1 for associated model information.

All other target plant responses showed no effect of *Epichloë* presence-related allelopathy, i.e., their 95% confidence intervals encompassed zero. Despite the positive allelopathic effect on shoot biomass, neither total nor root biomass showed a response (Figure 2A). Similar to shoot biomass, total and root biomass had statistically significant heterogeneity (total: Q119 = 204.31, *P* < 0.001; root: Q120 = 176.28, *P* < 0.001), with their I^2^ values suggesting low (root biomass) to moderate (total biomass) importance of effect size heterogeneity.^72^ Target plant floral biomass, number of leaves per plant, %-germination or -emergence, and nodulation score all had nonsignificant effect size heterogeneity, although the I^2^ value for leaves/plant (40%) suggests that heterogeneity might still be considered low to moderate. In contrast, shoot and root length had significant moderate to substantial effect size heterogeneity (Q49 = 87.22, *P* < 0.001, I^2^ = 41% and Q38 = 111.00, *P* < 0.001, I^2^ = 67%, respectively). For small-k responses (Figure 2B), the standard and adjusted random effects models differed in their detection of allelopathic effects for total arbuscule + vesicle numbers and vesicle numbers. However, fail-safe numbers were very low, and the individual studies showed no consistency in the direction of effects, suggesting that these models are not robust. Effect sizes for arbuscule numbers were highly heterogeneous.

None of the study design moderators were significant explanatory covariates of shoot biomass effect size (meta-regressions, Figure 2A). The lack of significance may be due to small sample sizes for some subgroups. However, small to moderate positive allelopathic responses were found for legumes, target plants treated with dried plant tissue, target plants in sterilized substrate, and those treated with substances from the *L. perenne*–*E. festucae* var. *lolii* host–endophyte combination. Application method and soil status were significant moderators of target plant root biomass responses to allelopathy (Figure 2B), with small and moderate positive responses for target plants treated with dried plant tissue and those in sterilized substrate, respectively. Neither target plant functional group nor the host–endophyte combination were significant moderators of target plant root biomass. Total biomass responded similarly to root biomass, although only the soil status moderator was statistically significant (see Online Resource 1). Finally, none of the moderators were significant explanatory covariates of shoot or root length, and none of the subgroup effect sizes differed from zero (Online Resource 1). Significant residual heterogeneity remained after accounting for moderators for all response variables.

## 4 Discussion

In our experiment and meta-analytical synthesis, we focused specifically on allelopathic effects due to the presence of the mutualist *Epichloë* endophyte in its grass host. Contrary to what has generally been hypothesized for these host-specific symbionts, we found very little evidence overall for negative allelopathic effects on other plants across a range of methodologies, target plant species, and response measures. Significant negative allelopathic effects were detected only for root hair measures, whereas positive effects were found for shoot, root, and total biomass in some experimental subgroups. The majority of response variables showed no evidence for allelopathy due to the presence of *Epichloë*. We also detected little allelopathy of, and few differences among, *Epichloë festucae* var. *lolii* strains.

The negative allelopathic effects detected for root hair density and length are based on multiple effect sizes from one study examining one legume genus (*Trifolium spp.*), using seeds that were grown for seven days on filter paper. ^66^ Thus, we do not know whether this result is replicable. Given that root hairs are involved in water and nutrient uptake and are the site of rhizobial infection and nodule formation in legumes,^73^ their reduction could negatively affect subsequent plant growth. However, the lack of allelopathic effects for root nodulation and legume root biomass and length, and the positive effect for legume shoot biomass, suggest that either Springer’s findings^66^ are anomalous or early changes to root hairs have negligible effects on subsequent plant growth. Microscopic examination for damage or changes at the cellular level may better indicate the mechanisms and consequences of the potential interaction.^5^

The overall positive effects on shoot biomass found for legumes and the *L. perenne*–*E. festucae* var. *lolii* pair suggest that *Epichloë* allelopathy is an unlikely mechanism explaining observations of poorer clover growth in endophyte-infected than uninfected *L. perenne* pastures. Experiments have shown that T. repens persistence in mixed pastures with *L. perenne* is affected by water availability,^27,74^ nitrogen fertilization level, ^75,76^ and grazing or mowing regime, ^77,78^ which influence competitive ability. Additionally, the *Epichloë* symbiosis can have various ecosystem effects ^79^ by altering resource competition,^80^ herbivore and pathogen pressure,^81^ and decomposition rates.^82,83; but^ ^see^ ^84^ These differences likely influence host-grass competitive outcomes in plant communities.

*Epichloë* presence in host grasses may alter soil microbial and invertebrate community composition as well as colonization of the host grass by arbuscular mycorrhizal and dark septate fungi, although studies do not always agree on the direction of effects. ^83,85–90^ Although some studies have suggested the possibility of negative plant-soil feedbacks to the infected host grass due to *Epichloë*,^68,91^ there were too few samples from the literature to determine whether *Epichloë* allelopathy might be involved (selfE+: *N* = 4 and 3 for shoot and root biomass, respectively); such allelopathy does not appear to affect the endophyte-free host grass. Rojas *et al*.^92^ noted that reductions in certain soil fungal genera with *Epichloë* presence in the grass host might indicate protective effects against plant pathogens, which could also benefit neighbouring plants. Soil microbes might also alleviate negative effects by metabolizing or transforming allelopathic substances. ^93^ One then might expect to see a more positive *Epichloë* allelopathic effect on target plant biomass in live soils than in sterilized soils. However, our meta-analysis detected a positive effect in sterilized soils and no effect in live soils for shoot and root biomass. Experiments testing the functional belowground effects of *Epichloë* presence ^e.g., 94^ may be fruitful in clarifying how the endophyte affects neighbouring plant performance via plant-soil feedbacks.^39,95^ Currently, there are insufficient measurements to draw conclusions about how *Epichloë* allelopathy affects arbuscular mycorrhizal fungal colonization of non-host plants, and there are no measures of effects on other microbes.

Although metabolite profiles of *L. perenne* root exudates differ among the endophyte strains we used in the greenhouse experiment,^50^ strains neither differed in allelopathic effects nor had a detectable effect on target plant biomass overall. Although we detected different effects for the legume *Glycyrrhiza lepidota*, and the direction was positive for two strains (AR1 and NEA2) compared to endophyte-free root exudates, the result is likely not biologically significant due to poor growth of this species during the experiment, possibly because it lacked its rhizobial symbiont. Limited data from the literature prevented meta-analytical comparisons among endophyte strains and among host plant genetic backgrounds (cultivars). We note that studies comparing *Epichloë coenophiala* strains in *S. arundinaceus* found few differential effects on potential sources of indirect plant-soil feedbacks: soil nitrogen fixation,^90^ arbuscular mycorrhizal and dark septate fungal colonization of the host grass, ^89^ or soil nutrients and function, with the exception of carbon. ^49^

It is difficult to evaluate adequately the true effect of bias in meta-analyses, particularly when effect sizes are heterogeneous and non-independent.^63,64,96^ One source of bias occurs when nonsignificant results are not published (i.e., publication bias or selective reporting). In our case, we were unable to obtain data from a few early studies, but we were able to obtain some unpublished data. Given the minimal number of significant effects in our meta-analysis, we do not think selective reporting was a major issue. In contrast, research bias could be problematic. For example, we only found *Epichloë* allelopathy studies for five host-endophyte pairs, all of which are well-studied agronomic grasses. In contrast, the most recent *Epichloë* taxonomic revision recognizes 29 distinct lineages (species, subspecies, and varieties) with strictly vertical transmission and 14 lineages with horizontal or both modes of transmission.^97,98^ Inferences from our meta-analysis may not extend to non-agronomic or horizontally transmitted *Epichloë* endophytes. Additionally, effect size data for many potentially important response measures were limited (e.g., nodulation, arbuscular mycorrhizal colonization, floral biomass, root hair characteristics) or nonexistent (e.g., seed production, survival, fitness, microbial activity).

The absence of *Epichloë* allelopathy contrasts the weight of evidence for allelopathy in grassland species, reviewed by Silva *et al*.^13^ where 73% of studies reported inhibitory allelopathic effects, and a further ∼22% reported both inhibitory and stimulatory effects. see also ^38^ Thus, the host grass species may have allelopathic effects on neighbouring plants, regardless of *Epichloë* presence, which we did not examine. Hodge and Fitter^99^, p. 867 note that the interpretation of plant-soil feedback studies “requires caution” because experimental conditions can be somewhat artificial, lacking community context, and potentially affecting the persistence and activity of allelochemicals. ^5^ Nevertheless, current evidence suggests that *Epichloë* allelopathy is unlikely to be a source of strong influence for suppressing neighbouring plant species or facilitating invasion of the host grasses into novel ecosystems.

**Figure 3:**
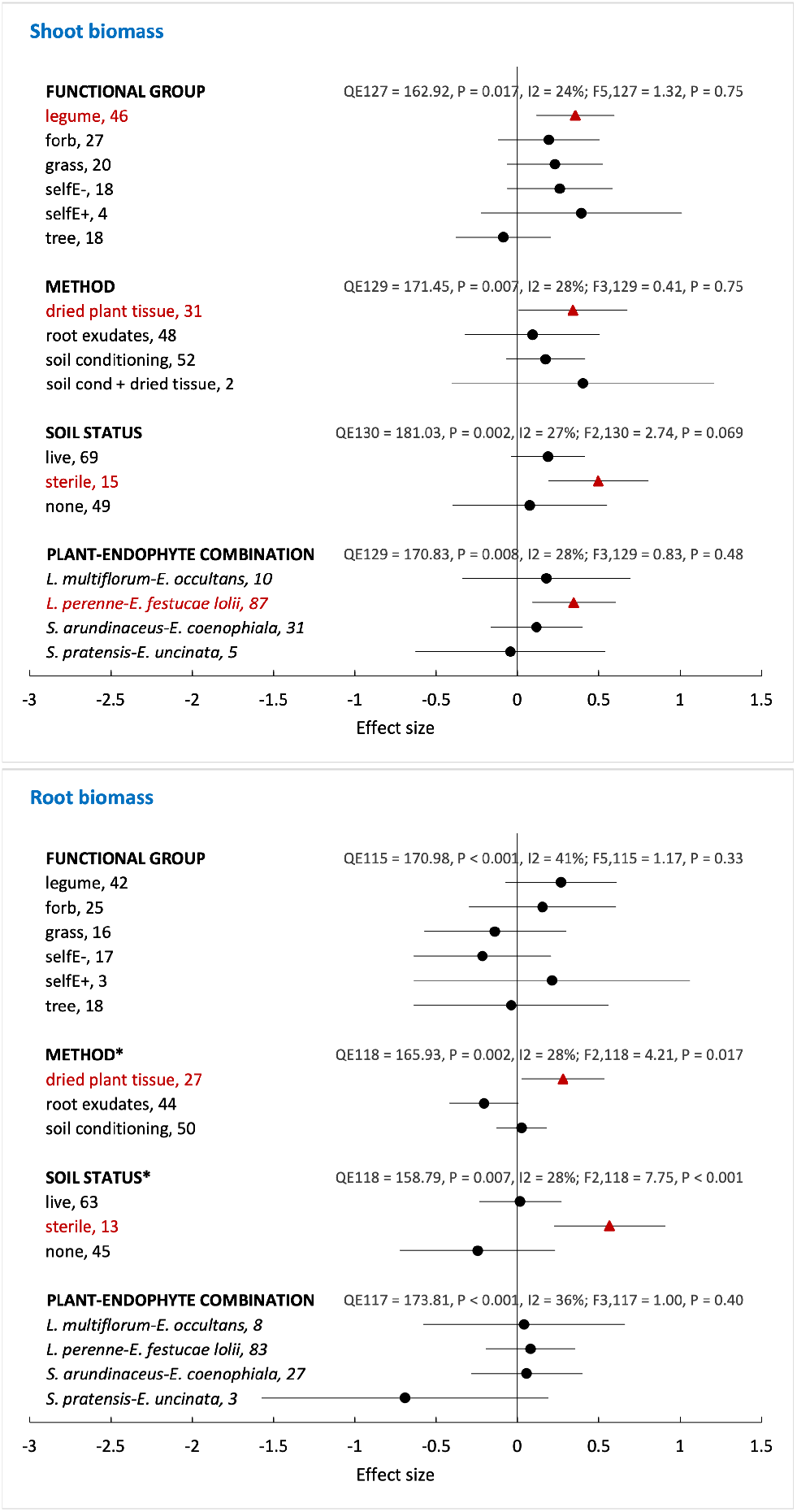
Effects of four moderators on target-plant responses to allelopathy due to the presence of *Epichloë* endophyte in its host grass for shoot biomass (A) and root biomass (B). Moderator test statistics are residual heterogeneity (QE) and its significance, proportion of heterogeneity not attributable to sampling error (I^2^), and omnibus *F*-test of moderator effect and its significance. Asterisk indicates significant moderator effect based on the omnibus test. Subgroup labels are followed by numbers of effect sizes. Points and whiskers are effect sizes and 95% confidence intervals, respectively. Dark red triangles = significant effects, black circles = nonsignificant effects

## Acknowledgements

This research was conducted on the ancestral lands of the Neutral, Anishnaabe, and Haudenosaunee peoples, and more recently, on the treaty lands and territory of the Mississaugas of the Credit First Nation. We recognize that today this gathering place is home to many First Nations, Inuit, and Métis peoples, and acknowledging them reminds us of our collective responsibility to the land where we learn and work, and to the on-going efforts for reconciliation. We gratefully acknowledge the assistance of N. Charlton, Z. Gedalof, several student volunteers, and the University of Guelph Phytotron staff. We thank C. Inch for providing the Lolium perenne seed and A. Patchett for hydroponics design. We also thank the authors who directly provided us their raw or summarized data for the meta-analysis.

## Data availability

The complete experimental and meta-analytical data sets are archived at: Hager HA, Gailis M, Newman JA (2021) Allelopathic effects of Epichloë fungal endophytes: experiment and meta-analysis. [data set] Scholars Portal Dataverse. https://doi.org/10.5683/SP3/VF5XWU

## Author contributions

Designed the experiment: HAH; performed greenhouse experiment: MG, HAH; performed meta-analysis: HAH; analysed data: HAH; wrote manuscript: HAH; edited manuscript: HAH, JAN, MG

